# Release of extracellular matrix components after human traumatic brain injury

**DOI:** 10.1101/2023.02.23.529754

**Authors:** Michael Bambrick, Mark Johnson, Jeffrey D. Esko, Biswa Choudhury, Alejandro Gomez Toledo, Kevin Staley, Ann-Christine Duhaime

## Abstract

Most research on the evolution of damage after traumatic brain injury (TBI) focuses on cellular effects, but the analysis of human tissue slices and animal research have shown that TBI causes concomitant damage in the extracellular matrix, which can play a significant role in both short-term consequences such as edema, and late effects such as post-traumatic epilepsy (PTE). To test the hypothesis that traumatic brain injury (TBI) in human patients causes disruption of sulfated glycosaminoglycan (sGAG) in the extracellular matrix, we measured levels of these substances in the ventricular cerebrospinal fluid (CSF) in patients with severe TBI in the acute post-injury period, along with concomitant levels in blood and urine. We assessed whether levels corresponded to parenchymal injury load, distance of traumatic brain lesions from the ventricle, presence of polytrauma, or host demographic factors.

**Methods:** Samples of CSF, blood, and urine were obtained within 72 hours of injury in patients who received external ventricular drains as part of their treatment of severe TBI, and levels of chondroitin and heparan sGAGs were measured, along with their disaccharide constituents. Basic demographic information, presence and severity of polytrauma, brain injury load based on imaging findings, and distance of radiologically visible parenchymal injury from the ventricle were analyzed for correlation with total subtype sGAG levels in each patient.

**Results:** Levels were measured in 14 patients ranging in age from 17-90 years. CSF sGAG levels were variable among patients, and sGAG levels were higher in plasma than in CSF and variable in urine. Patients with polytrauma had non-significantly higher blood sGAG compared to patients with isolated head injury. Subcategories of CSF sGAG levels correlated with distance from the ventricle of parenchymal injury but not with brain injury load, which may reflect rapid metabolism in the parenchyma, contamination by blood, or bulk directional CSF flow from the ventricle to the subarachnoid space.

**Conclusion:** This study is the first to measure sGAG levels in ventricular CSF and also provides the first measurements in patients with TBI. Damage to the extracellular matrix may play a major role in acute and chronic injury sequelae, and these data demonstrate elevation locally of intracranial sGAGS after severe TBI and also suggest rapid local metabolism of these breakdown products. The consequences of extracellular matrix breakdown may provide unique therapeutic and preventive avenues to mitigate post-injury sequelae.

## Introduction

The extracellular matrix (ECM) is an important component of solid organs. In fact, by holding the component cells together, the ECM is what makes a solid organ solid ^1^. The ECM fills the extracellular space ^2^, which comprises approximately 20% of the brain’s volume ^3 4^. The ECM is a hydrogel that is filled with extracellular fluid, proteoglycans, which are glycoproteins that contain long, polymeric sulfated glycosaminoglycans (sGAG), and hyaluronan (hyaluronic acid), a non-sulfated GAG ^5^ (Figure 1). The glycosaminoglycans are assembled by the copolymerization of alternating residues of N-acetylglucosamine and glucuronic acid in heparan sulfate and hyaluronan (although in different linkages), N-acetylglucosamine and galactose in keratan sulfate, and N-acetylgalactosamine and glucuronic acid in chondroitin sulfate/dermatan sulfate. The chains undergo variable modifications including deacetylation of N-acetylglucosamine residues in heparan sulfate, O-sulfation at various positions, and epimerization of glucuronic acid to iduronic acid. The disaccharide constituents of the GAG are named based on a four-character code reflecting their specific structures, including numbers denoting their sulfation characteristics (disaccharide structure code, DSC) ^6^. In the brain, chondroitin sulfate and heparan sulfate are the most plentiful sulfated GAGs (sGAGs), and hyaluronan is the most plentiful nonsulfated GAG ^7^. Within these categories, GAGs are further characterized based on the specific disaccharide component characteristics, including the position and number of sulfate groups in sulfated GAGs ^6^. Disaccharide characteristics can vary among species and sites and during development and plasticity ^7^.

**Figure 1.**
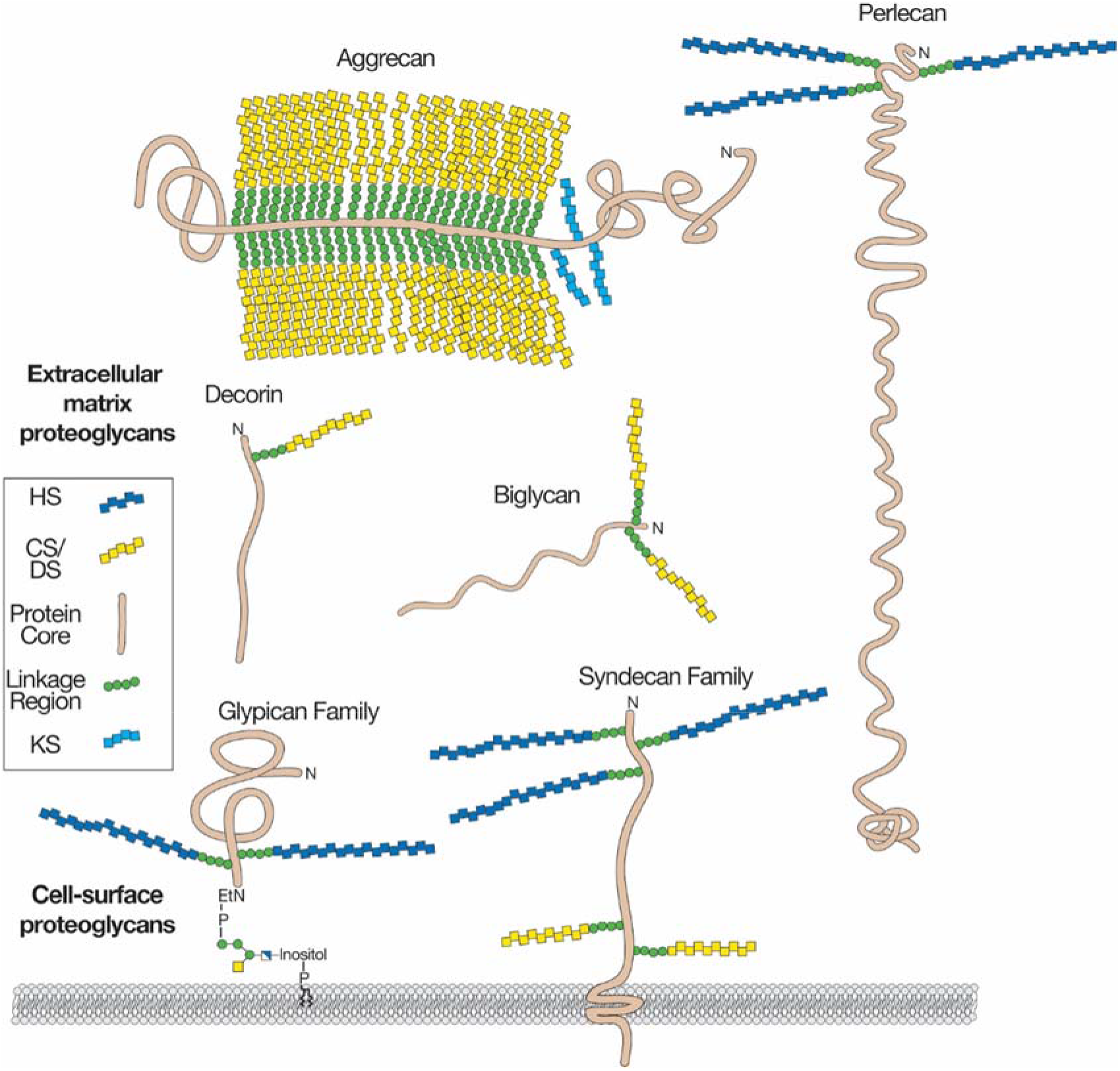
Proteoglycans consist of a protein core (brown) and one or more covalently attached glycosaminoglycan chains (dark blue, HS, heparan sulfate; yellow, CS/DS, chondroitin sulfate/dermatan sulfate; light blue, KS, keratan sulfate). Membrane proteoglycans either span the plasma membrane (type I membrane proteins) or are linked by a glycosylphosphatidylinositol (GPI) anchor. Extracellular matrix proteoglycans are usually secreted, but some proteoglycans can be proteolytically cleaved and shed from the cell surface (not shown). Reprinted with permission from Merry, C. L. R., Lindahl, U., Couchman, J., and Esko, J. D. (2022) Proteoglycans and Sulfated Glycosaminoglycans. In Essentials of Glycobiology (Varki, A., Cummings, R. D., Esko, J. D., Stanley, P., Hart, G. W., Aebi, M., Mohnen, D., Kinoshita, T., Packer, N. H., Prestegard, J. H., Schnaar, R. L., and Seeberger, P. H. eds.), 4th Ed., Cold Spring Harbor (NY). pp 217-232.

The sulfation of the disaccharides imparts a high negative charge density to the ECM ^8 9^. These anatomically fixed charges dramatically reduce the diffusion of charged molecules through the ECM by electrostatic interactions ^3 8^. We have proposed that, just as in the intracellular space ^10 11^, fixed charges in the extracellular space reduce the local concentration of extracellular chloride via Donnan exclusion ^12 13^. This creates unique local chloride concentration gradients across the neuronal membrane. In turn, these unique gradients may alter the direction and effects of activation of local inhibitory GABA_A_ synapses.

When neurons die, the ECM that surrounded them is replaced ^14 15^. After significant insults such as severe traumatic brain injury, large numbers of neurons die in a short period of time, and the metalloproteinases that break down the extracellular matrix are detectable in the CSF and blood ^16 17^. The resultant (and possibly also traumatic) dissolution of the ECM and its component sulfated disaccharides should result in large influxes of chloride salts and water into the extra- and intracellular spaces ^12^. These fluid shifts may contribute to acute morbidities of TBI such as cerebral edema, for which more effective prevention and therapies still are needed ^18^. Chronic increases in sGAGs could degrade inhibitory GABAA chloride currents and thereby contribute to post-traumatic epilepsy.

We currently know very little about dissolution and resorption of the brain’s ECM under physiological or pathological conditions. Matrix metalloproteinases that are released and activated in response to injury hydrolyze the core proteins of the ECM, releasing GAGs ^19^. Proteoglycans and GAGs are catabolized in a complex series of enzymatic reactions within lysosomes ^20^ or by extracellular enzymes, such as heparanases or hyaluronidases. However, when neurons die en masse acutely after TBI, the fate of GAGs and proteoglycan fragments has not been studied. Catabolism of released GAGs by surviving neuronal, glial and vascular cells may occur. However, after major injuries there may be more GAGs released than the remaining capacity to catabolize them. If so, then GAGs would be released to the cerebrospinal fluid and blood after TBI. GAGs content and composition have not been measured in ventricular cerebrospinal fluid (CSF), nor have they been studied as biomarkers after trauma ^21^. In this study, we measured GAG levels in CSF, blood, and urine in human patients treated with ventriculostomies for management of increased intracranial pressure after TBI. Our hypotheses were that CSF levels of GAGs would be increased compared to historical data derived from lumbar puncture, would increase with higher brain injury load or other injury characteristics, and that polytrauma patients might have higher blood and/or urine GAG levels compared to patients with isolated head injury.

## Methods

### Sample Collection

The data presented are a part of the Citizen’s United Research in Epilepsy’s (CURE) multi-center research study. Two Level 1 trauma centers, Massachusetts General Hospital and University of Massachusetts Medical Center, enrolled patients with severe acute traumatic brain injury (Glasgow Coma Scale less than or equal to 8) who required the placement of an external ventricular drain for clinical purposes. Ventriculostomies are placed at the treating team’s discretion, and in general are used in these two centers for patients with swelling-prone injury scenarios to manage anticipated or ongoing elevations in intracranial pressure ^22^. Patients with a pre-existing history of epilepsy or additional confounding neurologic conditions such as stroke, hydrocephalus, or neurodegenerative conditions were excluded.

The CSF and urine samples of the subjects all were collected as waste material, either discarded extra fluids obtained during clinical sampling, or discarded fluids during routine care when drainage bags were replaced. Blood samples were obtained from clinical lab excess material. Samples from urine and blood were timed to be obtained as closely as possible to the time of CSF sampling, typically simultaneously or within hours. No excess samples were drawn for the purpose of the study, and no direct contact with patients/families was made by the investigators, who coordinated obtaining excess samples directly from the treating team or clinical labs. As a result, the Institutional Review Board granted a Waiver of Consent for use of waste samples for the purpose of the study.

Samples of urine and CSF were frozen and stored for later sulfated GAG analysis, while the blood samples were centrifuged, and the separated plasma was extracted then subsequently frozen along with the other samples. These samples were obtained within the first 72 hours of injury to standardize the time window as much as possible within the limitations of the Waiver of Consent guidelines. To assess changes over time, additional samples were acquired during subsequent 72-hour time epoch windows if the functioning external ventricular drain was still in place.

The samples were analyzed at the University of California San Diego by the GlycoAnalytics Core (University of California San Diego, La Jolla, CA, USA). Samples were processed to analyze concentrations of sGAG subtypes chondroitin sulfate and heparan sulfate by digestion of the samples with chondroitinase ABC or heparin lyases and quantification of the disaccharide components as described below. CSF and urine (4 ml) were extracted, except for two of the samples in which lonly 2ml was available. GAG isolation from 200 μl of plasma samples were diluted using 200 μl of ice-cold DNase-RNase free Ultra-pure water. Then 375 μl of 2X-pronase digestion buffer (50 mM sodium acetate(NaOAc) with 200 mM NaCl, pH-6.0) was added and mixed thoroughly. Protease (25 μL (0.5mg), P5147 Sigma Aldrich) was added and the samples were incubated at 37°C for 16 h with vertical tumbling of the tubes.

### DEAE-Sephacel chromatography for isolation of GAG

Protease-treated samples were loaded on a small disposable chromatography column packed with 0.3 mL of prewashed dietheylaminoethyl (DEAE) Sephacel matrix. The pre-washing of the DEAE-Sephacel matrix was done using 2 ml of DEAE-equilibration buffer (50 mM NaOAc with 200 mM NaCl and 0.1% Triton X-100, pH-6.0). Samples were loaded on the column and allowed to flow under gravity. GAG molecules are negatively charged and bind to the DEAE-resin through charge interaction. The sample tube was rinsed with 1 ml of DEAE-wash buffer to ensure complete transfer of the samples. DEAE-column was washed with 6 ml (20x bed volume) of DEAE wash buffer and finally the bound GAGs were eluted in 2.5 ml of DEAE elution buffer (50 mM NaOAc with 2 M NaCl, pH-6.0) in a 15 ml tube. Samples (2.5 mL) were loaded on conditioned PD10 desalting columns (10% ethanol/water) and samples were allowed to flow through completely. Finally, GAGs were eluted using 3.5 ml of 10% ethanol solution and lyophilized.

### Heparan Sulfate and Chondroitin Sulfate analysis

Lyophilized GAG underwent chondroitin sulfate (CS) and heparan sulfate (HS) digestion using chondroitinase ABC and heparinase I, II and III, respectively. After digestion the samples were purified using 3K spin filtration, tagged with [^12^C]aniline, mixed with [^13^C]aniline tagged disaccharide standards and analyzed by liquid chromatography/mass spectrometry ^23^. We analyzed the distribution of component disaccharide species in these samples in order to assess the degree to which the released GAGs were sulfated, which might contribute to ionic shifts with physiologic consequences. The values for the individual disaccharides were summed to determine the total amount of GAG recovered in each sample.

### CT/MRI Imaging Analysis

Following collection of the samples, the subjects’ clinical imaging results and images were reviewed by a neurosurgeon with background in standardized neurotrauma image analysis (ACD) blinded to GAG levels to determine both the imaging-derived injury load and the distance of the closest visually damaged area to the lateral ventricles where the EVD was placed. The brain *injury load* was scored based on a summation of 5 different scores assigned to each patient (total range 0-21 points), using the National Institutes of Health Common Data Elements definitions for different types of traumatic lesions ^24^. Extradural injury (skull fracture or hemorrhage) was scored as 0 if absent and 1 if present. Supratentorial parenchymal hemorrhage/contusion was scored as present or absent in each lobe bilaterally, with scores thus ranging from 0 to 8. Subdural and/or subarachnoid hemorrhages were scored similarly by lobe, with possible scores also ranging from 0-8. Posterior fossa injury was scored as 0 if absent, 1 if parenchymal injury or subdural/subarachnoid hemorrhage was 1-2 cm in diameter, and a score of 2 if larger areas were involved. An additional point was added for gross intraventricular hemorrhage (IVH larger than 5mm), and another point for midline shift measured at the foramen of Monro greater than or equal to 1cm. Thus, using this system, total injury load score had allowable values from 0-21. In addition to imaging load, the distance from the nearest parenchymal contusion/hemorrhage site to the lateral ventricle was determined by measuring from the edge of the lesion to the edge of the ventricle in mm. Brain injury load and distances were corroborated by a second, independent reviewer (MJB). All images analyzed were obtained within the first 72 hours of injury, occurring within the initial time epoch of sample collection.

### Trauma Classification and Scoring

Subjects were classified as having isolated head trauma or multi-system trauma by records review. Patients with involvement of head only or head plus a single low-severity extremity injury were classified as predominantly isolated head injury, while those with more severe and multi-region injuries were classified as multi-system trauma. Injury body region distribution and severity were verified using the AIS (Abbreviated Injury Score) and ISS (Injury Severity Score) for each individual when available; these scores sum injury subscores for each body region. The severity of extracranial trauma was quantified by summing the AIS subscores and subtracting the score for head injury. The ISS is a more detailed score reflecting overall injury severity, including of the head, and was utilized for correlation with measured GAG values.

### Statistical Analysis

Differences among levels of different disaccharides were determined by ANOVA test and subsequent post-hoc analysis to assess for significance of differences. Injury-related variables, including severity of extracranial trauma (AIS score values excluding the AIS head score), ISS, imaging load, and proximity to ventricle (as previously described), and age as a demographic variable were presented as continuous variables. The correlation of individual sGAG results to injury-related variables and age were assessed using a Pearson correlation coefficient equation, and subsequently calculated t value. These were then used to calculate the p value and it was adjusted to reflect the number of independent tests performed. While there are no normative values for ventricular CSF GAGs from healthy patients, levels also were compared to those obtained from lumbar puncture in healthy controls collected for other purposes, acknowledging potential confounding differences from these sources (discussed further below) ^25 26 27^.

## Results

Patients ranged in age from 17-90 years of age. Mean time from injury to initial CSF sampling was 68 hours and 51 minutes (standard deviation of 49 hours and 11 minutes). Four patients also had samples in the 4-6 days post-injury epoch, and one patient had a third set of samples at 9 days after injury. Of 10 patients with full extracranial injury profiles, 4 had isolated TBI and 6 had polytrauma. Of the polytrauma patients, extracranial injury severity (AIS minus head injury severity scores) ranged from 3-19 (mean 9.8), reflecting a severity range from mild to critical thoracoabdominal, skeletal, and soft tissue injuries.

Collection of CSF from ventricular catheters/drainage reservoirs, as well as collection of blood and urine samples from excess laboratory or drainage samples, was readily accomplished with no risk to patients at both institutions. No complications or limitations of these procedures were found. Storage, shipment, and remote analysis of samples were accomplished without losses. Thus the study demonstrated that multicenter collection and analysis of ventricular CSF, blood, and urine for GAGs after severe head trauma could be feasibly executed as a minimal risk prospective human trial in patients with traumatic coma.

For this report, we analyzed only the initial time point of sampling for each patient, within the first 72 hours of injury. Of the 14 patients with samples analyzed, a total of 21 unique CSF sGAG levels were analyzed (2 subjects had the values from a single time point averaged due to multiple tests run on the same sample).

### Chondroitin sulfate in ventricular CSF

Of n=21 samples run, the mean concentration of sulfated CS GAGS in the CSF was 218 ng/ml ± 224 (SD). These results fall below the range for normative mean levels in pediatric patients from lumbar CSF samples, which had a mean of 670 ng/ml ± 570 ng/ml ^25^. The individual disaccharides present in the analysis showed a nonuniform distribution, with D0a4/D2a0, a disaccharide with a single sulfate group, having the greatest concentration, with a mean of 149 ± 152 ng/ml. Analysis using ANOVA with a post-hoc test showed that this disaccharide had significantly greater concentration than the rest of the disaccharides measured in the study, with a p value that was < 0.005 in relation to the other disaccharides. Among the other disaccharides, D0a0, which is not sulfated, had a mean concentration of 28 ± 36 ng/ml. D0a6, which has one sulfate group, was 26 ± 34 ng/ml. D2a4, which has two sulfate groups, was 2.6 ± 3 ng/ml. D2a6, also having two sulfate groups, was 2.6 ± 3 ng/ml. D0a10, with two sulfate groups, was 9.6 ± 9 ng/ml. Finally, D2a10, with three sulfate groups, was 0.03 ± 0.05 ng/ml. D0a4/D2a0 was found in the largest concentration, with D0a0 and D0a6 having the next closest grouping. D0a10 also had a concentration greater than D2a4, D2a6 and D2a10. The latter three all had relatively low concentrations. In total the mean concentration of the sulfated disaccharides was 190 ng/ml while the nonsulfated group was 28 ng/ml.

**GRAPH 1:**
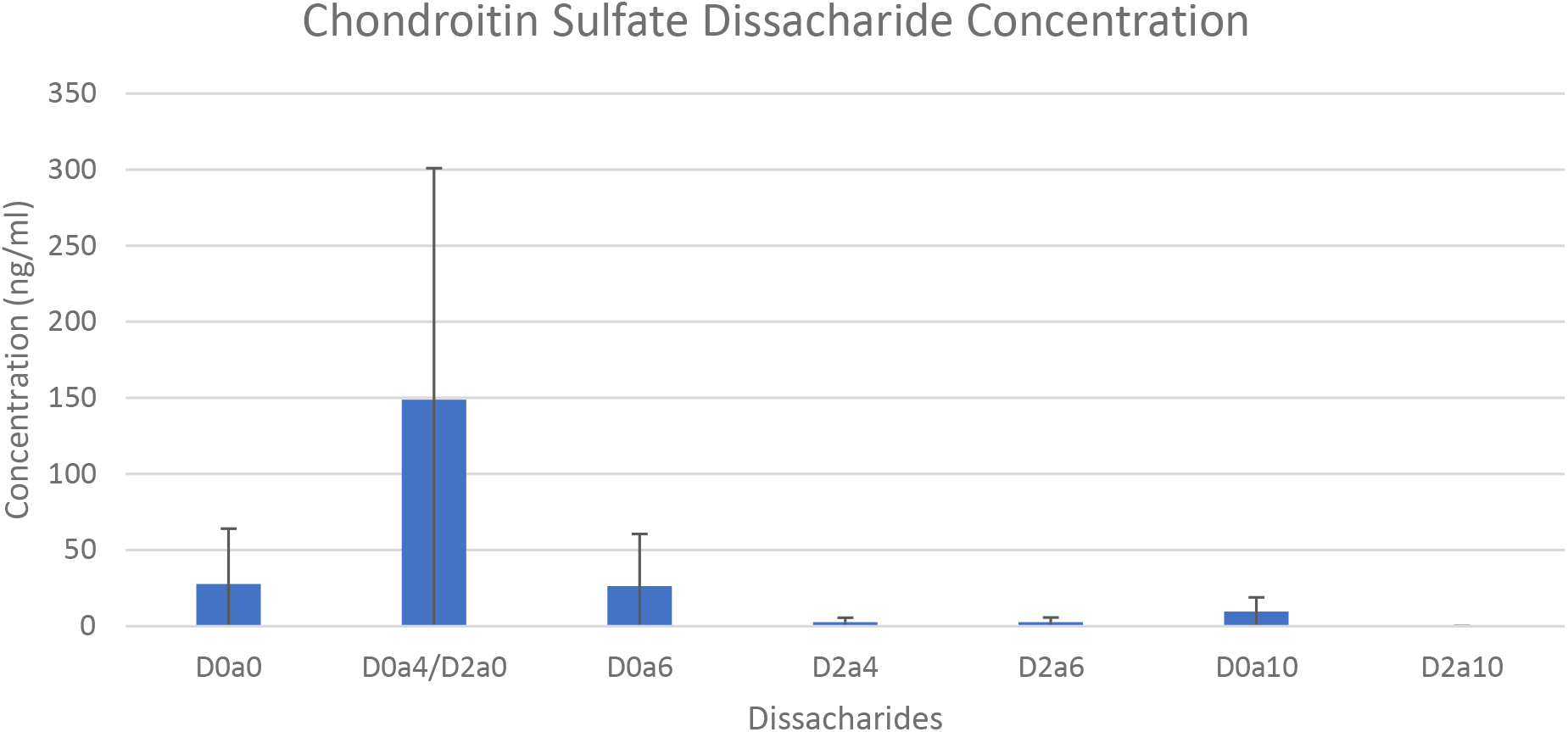
Concentration of chondroitin sulfate disaccharides within cerebrospinal fluid samples. Concentration is measured in ng/ml with D0a4/D2a0 disaccharide showing the highest average concentration and a statistically significant difference from the rest of the disaccharides (p<0.005).

### Heparan sulfate in ventricular CSF

The CSF samples digested by heparinase I, II and III (n=21) showed a mean concentration of HS GAGs of 23 ± 17 ng/ml, which is a low value when compared to the normative results from lumbar punctures of 50 ± 16 ng/ml ^26^. Similar to the CS sGAGs, a single disaccharide, D0A0, which is not sulfated, showed the greatest concentration at 12 ± 10 ng/ml. Performing an ANOVA and post-hoc test on the data set showed that the D0A0 disaccharide had statistically significant differences from all of the rest of the disaccharides, with a p value < 0.0005 for all comparisons. For the additional disaccharides, the concentration of D0H0, which also is not sulfated, was 0.04 ± 0.09 ng/ml. For the disaccharides with one sulfate group, D0H6 had a concentration of 0.002 ± 0.004 ng/ml, D2H0 was 0.001 ± 0.003 ng/ml, D0S0 was 3.4 ± 2.9 ng/ml, D0A6 was 1.8 ± 1.3 ng/ml and D2A0 had a concentration of 0.18 ± 0.15 ng/ml. For the species with two sulfated groups, D2H6 had a concentration of 0.074 ± 0.13 ng/ml, D0S6 had 1.1 ± 4 ng/ml, D2S0 had 2.2 ± 1.6 ng/ml, and D2A6 had 0.002 ± 0.005. Finally, D2S3, which possesses three sulfated groups, had a concentration of 1.1 ± 0.7 ng/ml. In total, the average concentration of the sulfated disaccharides was 8.2 ng/ml, while the disaccharides without a sulfate group had a concentration of 11 ng/ml. This contrasts with the results in CSF digested by chondroitinase ABC, which had a higher concentration of sGAGs than non-sGAGs.

**GRAPH 2:**
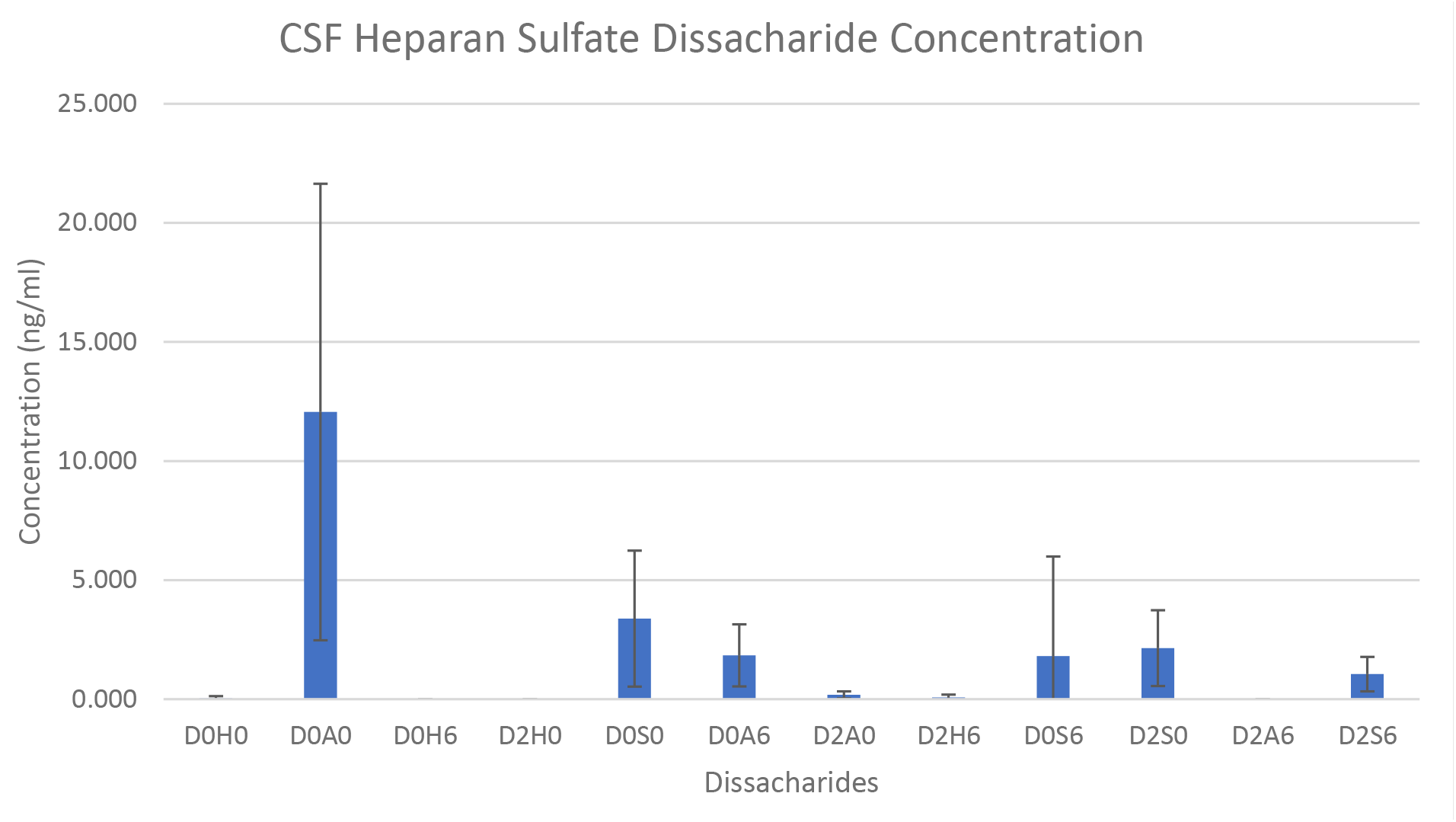
Concentration of heparan sulfate disaccharides in CSF samples. Concentrations are measured in ng/ml with the D0A0 disaccharide having the highest mean concentration and statistically significant value compared to the rest of the disaccharides (p<0.0005).

### Plasma and Urine sGAG levels

The levels of sGAGs found in both CS and HS plasma samples were greater on average than those found in the ventricular CSF. There were 18 unique plasma samples obtained, with some larger samples for a single time point being analyzed separately and then subsequently averaged. Plasma CS sGAGs were found to have a concentration of 7.4 μg/ml, which is an order of magnitude larger than the concentration found in the CSF. For HS, the concentration of sGAGs was found to be 0.23 μg/ml, which is also an order of magnitude larger than that in CSF samples. For assessment of urine, there were a total of 12 unique samples, reflecting Waiver of Consent constraints that did not allow for clinically unnecessarily sample collection during every time epoch. The levels found in the urine samples were found to have a concentration of 3.1 μg/ml for CS, and 1.7 μg/ml for HS, also greater than CSF levels.

### CSF and Plasma sGAG Concentration Comparison

Comparing the levels of sGAGs in CSF and plasma highlight a difference in the concentrations. To further elucidate this difference and test whether a relationship is present, the concentrations were plotted against one another and separated into groups of samples from patients with isolated head trauma compared to those with multitrauma. With CS, the Pearson correlation coefficient between isolated head trauma CSF and plasma sGAG values was found to be −0.36. This was not statistically significant as the p value of the correlation coefficient was found to be 0.61. For multitrauma, the Pearson correlation coefficient of 0.18 also was found to not be statistically significant with a p value of 1.23. These data suggest no statistically significant correlation between CSF and plasma sGAG levels in either multitrauma or isolated head trauma in the CS samples.

**GRAPH 3:**
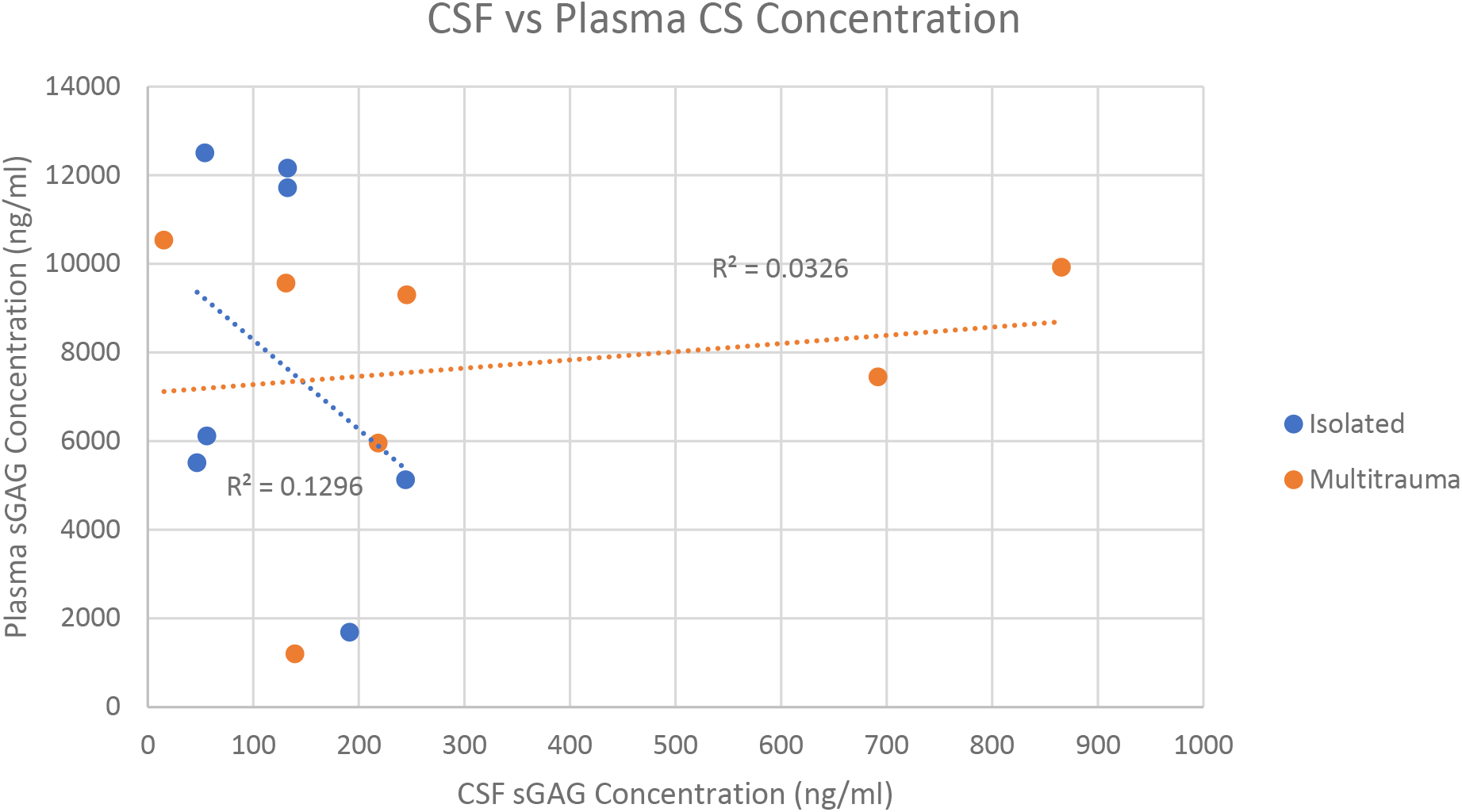
Plasma and CSF sGAG concentrations (ng/ml) of chondroitin sulfate plotted against one another and separated into groups denoting isolated head trauma or multitrauma. No correlation was observed either within the groups or between the groups. The linear regression of each of the individual groups was also not found to be significant.

For the HS samples, there also was no significant relationship between CSF concentrations and blood concentrations. The Pearson correlation coefficient for isolated head trauma was −0.35 with a p value of 0.63 and for multitrauma the coefficient was −0.58, with a p value of 0.15.

**GRAPH 4:**
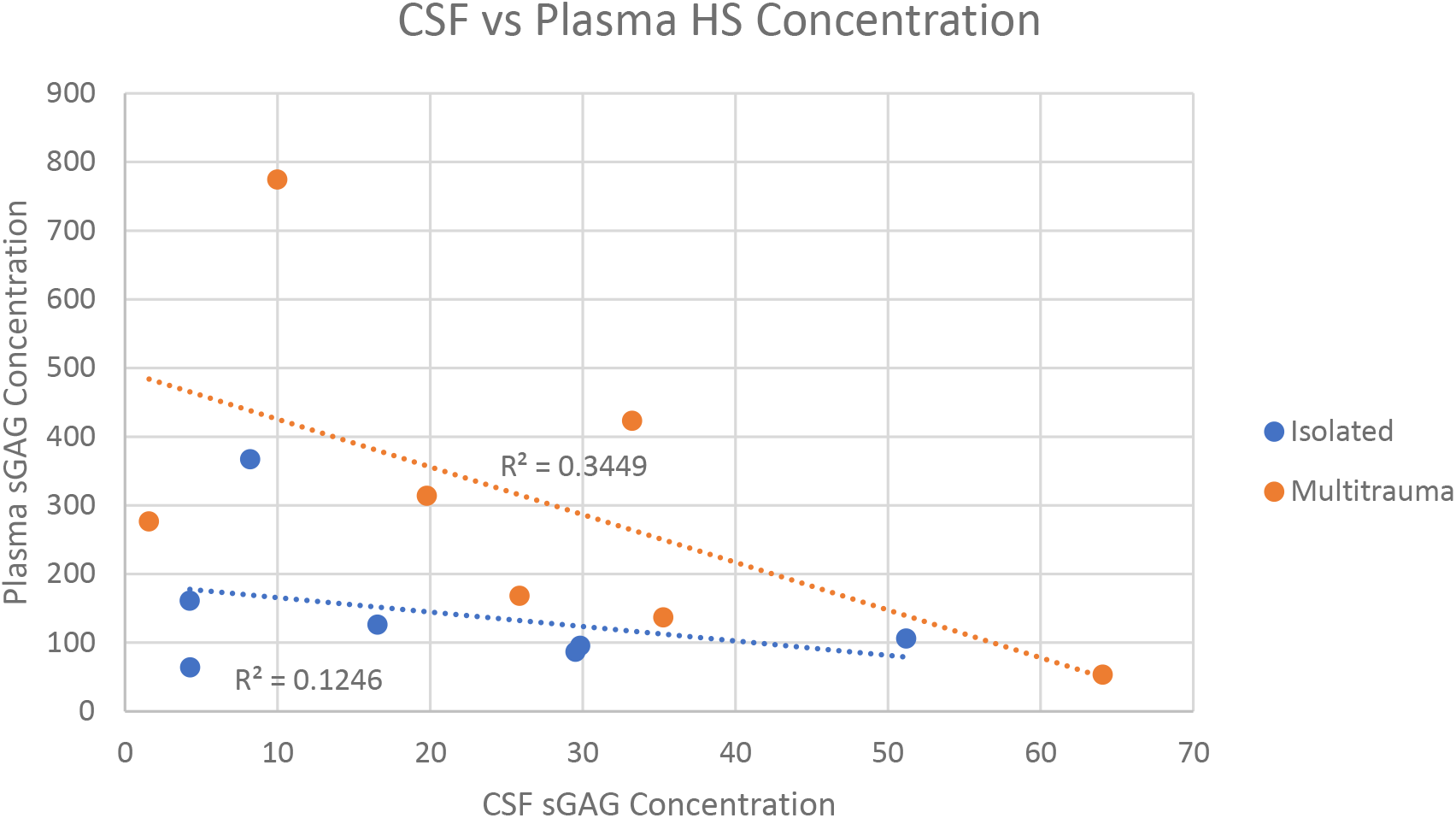
Plasma and CSF sGAG concentrations (ng/ml) of heparan sulfate plotted against one another and separated into groups denoting isolated head trauma or multitrauma. No correlation was observed either within the groups or between the groups. The linear regression of each of the individual groups was also not found to be significant. While plasma levels on average were higher in multitrauma patients compared to those with isolated head injury, these differences were not significant.

### Trauma Dependent Effects

In both CS and HS sGAG concentrations, individual subjects displayed a wide range of values. In order to assess if there is a trauma-dependent effect the values were assessed against extracranial trauma, imaging load, ISS, distance from lateral ventricle, and age.

There was no correlation between the concentrations CSF CS sGAGs to degree of extracranial trauma, ISS, radiologic assessment of cerebral injury load, distance of injury to lateral ventricle, or patient age. Proximity of radiologically visible traumatic brain injury to the lateral ventricle had the strongest correlation, though still not significant with a p value of 0.55. Similarly, no correlation was found between extracranial trauma, ISS, imaging load, or age and CSF HS sGAG concentrations. However, the concentration of HS sGAGs in the ventricular CSF was found to correlate with the distance of the visible injury from the lateral ventricle. The Pearson correlation was found to be −0.75 with a p value of 0.048.

**GRAPH 5:**
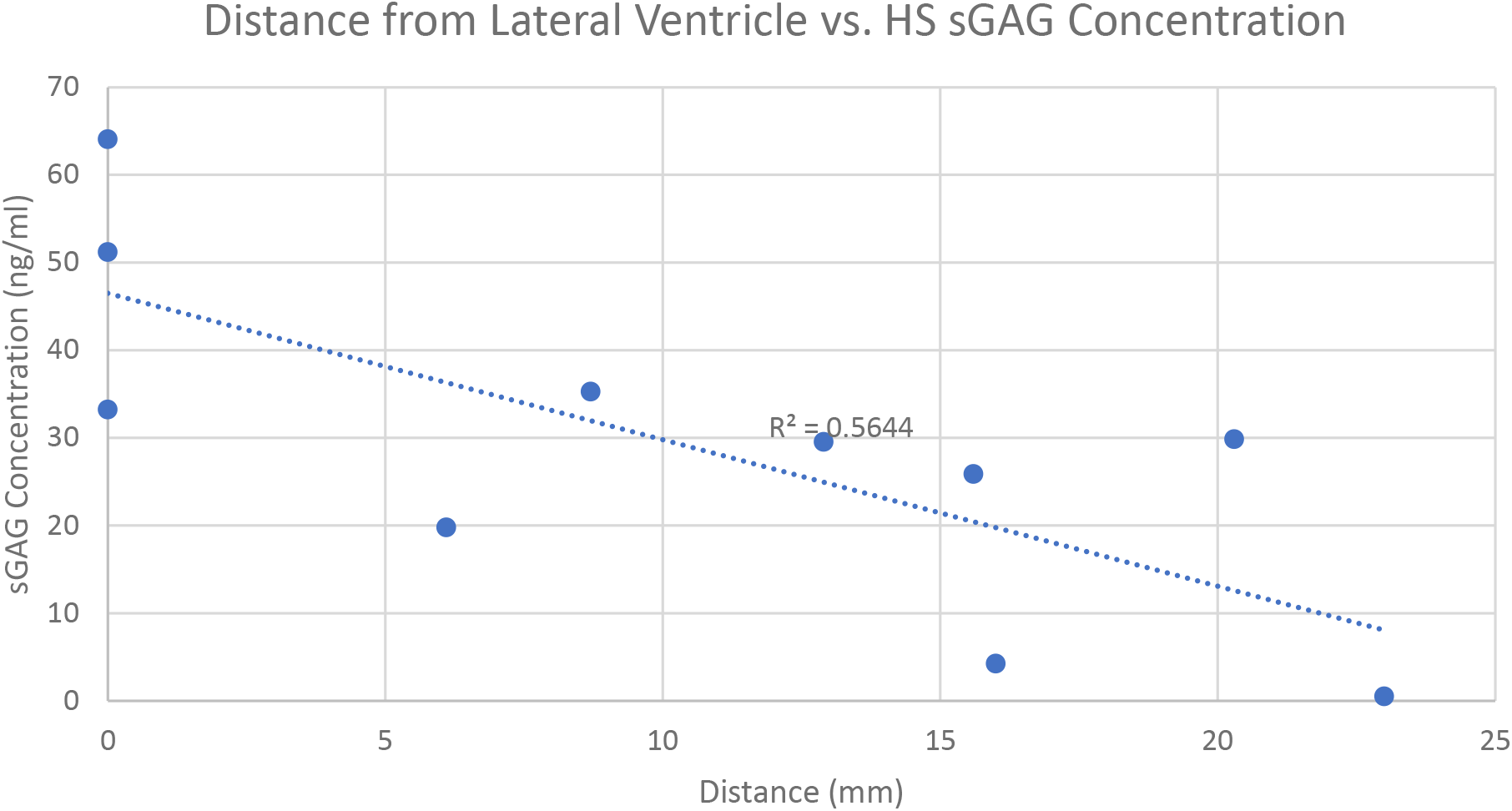
Heparan sulfate concentration in CSF samples plotted against distance from the lateral ventricle where the external ventricular drain was placed. The Pearson correlation was found to be −0.75 with a p value of 0.048. This effect was not found in chondroitin sulfate CSF samples.

## Discussion

This study describes sGAG levels in the ventricular CSF, blood, and urine of human patients with severe head trauma requiring ventricular CSF drainage for management of intracranial pressure. Feasibility was demonstrated for the minimally invasive sampling of CSF (in the setting of an indwelling ventricular catheter for clinical care), blood, and urine; multicenter collaboration; and remote analysis of GAGs at a specialized center. The levels of CSF sGAGs were low relative to prior analyses of normal human lumbar CSF. There were borderline correlations between ventricular GAG levels and the proximity of traumatic lesions to the ventricles. The small sample size, the inherent variability of traumatic brain injury, and the natural variation in sGAG content and composition across individuals may have limited the significance of the findings.

To the authors’ knowledge, this is the first report of GAG concentrations in human ventricular CSF. Ventricular CSF is not available from control patients due to the necessarily invasive nature of the sampling procedure. These samples were obtained from comatose trauma patients whose ventricular CSF was being drained to manage increased intracranial pressure. Despite the severity of the brain injury, the ventricular CSF GAG levels were lower than human control CSF obtained from lumbar puncture ^25 26 27^. Below we consider two potential explanations for this finding.

Lumbar CSF has higher levels of protein and cells than ventricular CSF ^28^. This is thought to be a consequence of the proximity of ventricular CSF to the choroid plexus, where CSF is produced. In contrast, CSF collected from the lumbar space has diffused over the surface of the cortex and brain, providing a greater opportunity to pick up protein and cells extruded from these surfaces. The same may be true for GAGs. Experimental studies of cerebral edema have found that neutral and positively charged molecules can flow into ventricular CSF ^29^, but molecules that are negatively charged at physiological pH, such as albumin and fluorescein isothiocyanate, flow to the subarachnoid CSF ^30^. The negatively charged sGAGs therefore may not have been cleared via a directional pathway into ventricular CSF.

Another potential explanation for the low levels of ventricular CSF sGAGs in trauma patients is that sGAGs are not released into any CSF space under these conditions. In other words, after severe brain injury, dissolution of the extracellular matrix ^31^ may not result in significant spillage of matrix components into either the ventricular or subarachnoid CSF. Most of what is known regarding the catabolism of GAGs is based on studies of genetic defects in GAG catabolism. These defects form the bases for the human mucopolysaccharidoses such as Hunter’s, Hurler’s, and Sanfilippo Diseases. In these disorders of GAG catabolism, neurons, glia, and vascular cells accumulate the partially degraded GAGs that cannot be further broken down due to enzyme deficiency ^32^. GAG components that are not catabolized are also stored in peripheral organs and excreted in the urine; however there is no evidence that these GAGs arise from the brain rather than the peripheral organs. Lumbar CSF levels of GAGs are modestly elevated in these disorders ^25 26^. Thus there is not much evidence for substantial spillage of partially catabolized GAGs *from the brain* into CSF, blood, and urine in genetic disorders of GAG catabolism. It is not known at what rate brain GAGs are released by matrix metalloproteinases after human trauma ^33 12^, nor the rate at which the released GAGs can be taken up by surviving neurons and glia. We speculate that after severe brain trauma, surviving neurons, glia, and perhaps vascular cells take up the majority of the GAGs released by matrix metalloproteinases, limiting GAG spillage into the CSF and bloodstream.

The modest correlations between HS and CS levels and proximity of traumatic lesions to the ventricular surface suggests that spillage of sGAGs into ventricular CSF occurs in at least some circumstances. This relationship bears further investigation, because in our cohort, at least some quantity of blood also was present in the ventricular CSF of patients with lesions that were closest to the ventricular surface. The blood may have had higher sGAG levels (Figures 3 and 4), resulting in spurious CSF sGAG elevation. Alternatively, the presence of blood and sGAGs in the ventricular CSF in patients with lesions near the ventricular surface may arise independently from damage to the periventricular ECM and the ependymal lining of the ventricles. Normally, the fixed negative charges borne by intact sGAGs in the periventricular ECM should limit the diffusion of anionic molecules by Donnan exclusion ^34 13^, and the flux of sGAGs should be further reduced by the ependymal lining of the ventricles ^35^.

This study was limited by the number of subjects, especially in light of the variability of both the traumatic injuries and sGAG levels. The study did not enroll for 6 months during the COVID-19 pandemic, and enrollment was limited thereafter by the reduction in motor vehicle usage and accident rates ^36^. In the absence of COVID, we would expect an approximate 50% increase in the rate enrollment over the course of this study. The study is also limited by a lack of controls; it is not ethical to obtain control human ventricular CSF. We focused on sGAGs, which are plentiful in the brain extracellular matrix, but there are many other matrix components that could also be studied ^5^. While this study eliminates one possibility for the fate of sGAGs released by matrix metalloproteinases after traumatic brain injury (transfer to CSF), it does not identify the path of further catabolism of the released sGAGs.

This pilot study will hopefully lay the groundwork for larger clinical studies by presenting the concentrations and distributions of the sGAGs and disaccharides present within the ventricular CSF and plasma in a two-center group of severely injured trauma patients. Addressing the mechanistic questions raised by this observational clinical study may best be accomplished using experimental preparations ^37^ in which both intra and extracellular sGAG levels can be analyzed after controlled subtypes of traumatic brain injury.

## Abbreviations

AIS: Abbreviated Injury Score
CS: chondroitin sulfate
CSF: cerebrospinal fluid
DSC: disaccharide structure code
CT: computerized tomography
DEAE: diethylaminoethyl
ECM: extracellular matrix
GABA: gamma aminobutyric acid
GAG: glycosaminoglycan
HS: heparan sulfate
ISS: Injury Severity Score
MRI: magnetic resonance imaging
PTE: post-traumatic epilepsy
sGAG: sulfated glycosaminoglycan
TBI: traumatic brain injury

